# Assessment of the hypoglycemic activity of trans-cinnamic acid isolated from the extract of *hairy rots Scutellaria baicalensis* in *in vivo* experiments

**DOI:** 10.1101/2024.09.12.612583

**Authors:** Fedorova Anastasia Mikhailovna, Milentyeva Irina Sergeevna, Asyakina Lyudmila Konstantinovna, Prosekov Alexander Yuryevich

**Affiliations:** Federal State-Funded Educational Institution of Higher Vocational Education “Kemerovo State University”

**Keywords:** trans-cinnamic acid, *Scutellaria baicalensis*, *hairy rots*, dietary supplements, *in vivo*, *in vitro*

## Abstract

Hypoglycemia is the process of reducing blood glucose concentration to below 2.5–2.8 mmol/L in men and less than 1.9–2.2 mmol/L in women. Prolonged hyperglycemia can lead to a range of serious chronic complications such as diabetic nephropathy, diabetic retinopathy, cardiovascular diseases in diabetes, and so forth, threatening human life, health, and safety. *Scutellaria baicalensis* is one of the most commonly used remedies in traditional Chinese medicine. This plant contains a number of biologically active compounds that contribute to improving kidney function, insulin resistance, and retinopathy in patients with type 2 diabetes, but few researchers study the toxic effects of medicinal plant components. Therefore, the aim of this study was to evaluate the hypoglycemic activity of trans-cinnamic acid isolated from the extract of *hairy rots of Scutellaria baicalensis* in *in vivo* experiments. This study involved intraperitoneal administration of alloxan at a dose of 150.0 mg/kg and oral administration of trans-cinnamic acid at doses of 50.0 and 100.0 mg/kg. During the experiment, it was found that trans-cinnamic acid at the presented doses does not affect the body weight dynamics of experimental animals with diabetes. Thus, this study demonstrates that trans-cinnamic acid at the presented dosage can be safely used as an ingredient in the creation of dietary supplements for the prevention of diabetes, thereby contributing to healthy aging.

## Introduction

The problem of combating diabetes becomes increasingly urgent and in demand from year to year. Lifestyle characteristics, consumption of high-carbohydrate, salty, fatty foods, synthetic additives, stressful situations, harmful habits, genetics—all contribute to the development of diabetes. The ecological situation in the world also influences the development of this pathology. Today, diabetes is one of the most common non-infectious diseases capable of leading to disability and death [1].

Despite the availability of many drugs on the pharmaceutical market for treating diabetes, active search for new therapeutic possibilities is underway. In many parts of the world, various dietary supplements (DS) occupy a significant portion of the pharmaceutical market, widely advertised in the mass media and used indiscriminately by many patients. The danger of the widespread use of DS lies in the fact that in some cases, patients take such supplements instead of conventional pharmaceuticals, leading to the development of severe complications in diabetes, including ketoacidosis [2].

Currently, before entering the market, DS do not undergo the established certification procedure regulated by special authorities. The dietary supplements industry is a self-regulating area devoid of necessary control [3].

An important issue with the use of dietary supplements is drug interactions. Patients often take DS indiscriminately, without paying attention to their composition, and freely combine them with each other and with other pharmaceutical drugs [4]. Therefore, the intake of DS should be coordinated with a physician who can assess potential drug interactions.

The ideal alternative therapy should have effectiveness similar to commonly accepted pharmaceuticals and lack their side effects. To achieve this, DS include substances derived from various plants, as well as vitamins and minerals.

Plants with hypoglycemic properties have been used in traditional medicine for a long time [5]. Many pharmaceuticals have originally been derived from plants. Specifically, one of the most widely used drugs for type 2 diabetes, metformin, was isolated from the flowering plant *Galega officinalis* (French lilac) [6]. However, in most cases, reliable information regarding the effectiveness, safety, and appropriateness of using herbs, vitamins, minerals, and other DS components in the treatment of diabetes is lacking. Considering the increasing prevalence of using such preparations without a physician’s prescription and supervision, it is important to assess the role of DS in the treatment of diabetes.

A number of plant substances with hypoglycemic effects are described in the literature; however, the study designs for most of them do not meet the principles of evidence-based medicine. For a more objective evaluation of the available data, referring to meta-analyses of database information is justified. Among the most studied plant components are the following: *Coccinia indica* (ivy gourd), *Panax ginseng* (Asian ginseng), *Ocimum sanctum* (holy basil), *Trigonella foenum-graecum* (fenugreek), *Ficus carica* (fig leaf), *Opuntia streptacantha* (Mexican cactus), and others [7, 8].

*Scutellaria baicalensis* (Baikal skullcap) is a medicinal plant containing a range of flavonoids (baicalin, baicalein, wogonoside, wogonin, etc.) that contribute to the comprehensive treatment of diabetes [9,10]. Currently, research on the effective components of *Scutellaria baicalensis* mainly focuses on flavonoids, while other components such as essential oil, amino acids, phenylethanol, and sterols are still insufficiently studied [11]. Presently, there are relatively few studies specifically dedicated to the toxic effects and pharmacokinetics of the active components of *Scutellaria baicalensis*. Therefore, it is necessary to continue large-scale experiments and further scientific research to fully assess the clinical efficacy and safety of this medicinal plant. Based on the above, the aim of this study is to evaluate the hypoglycemic activity of trans-cinnamic acid isolated from the extract of *hairy rots Scutellaria baicalensis* in *in vivo* experiments.

### Objects and methods of the study

The object of the study is trans-cinnamic acid, obtained from the aqueous-alcoholic extract of the root culture *in vitro Scutellaria baicalensis*.

Trans-cinnamic acid, isolated from the extract of the root culture of *Scutellaria baicalensis*, was obtained at earlier stages. The methods of cultivating the root culture of *Scutellaria baicalensis*, the extraction process, and the isolation and purification of trans-cinnamic acid are described in the work by A.M. Fedorova and her colleagues [12]. For the transformation of seedlings grown for 14–28 days on a nutrient medium with the following composition: macrosalts B5 – 50.00 mg, microsalts B5 – 10.00 mg, Fe-EDTA – 5.00 ml; thiamine – 10.00 mg; pyridoxine – 1.00 mg; nicotinic acid – 1.00 mg; sucrose – 30.00 g; inositol – 100.00 mg; 6-benzylaminopurine –0.05 mg; indoleacetic acid – 1.00 mg; agar – 20.00 g, soil bacteria strains *Agrobacterium rhizogenes* 15834 Swiss (Moscow, Russia) were used. For the *in vitro* root cultures of *Scutellaria baicalensis*, the cultivation cycle was 5 weeks. The parameters for obtaining the aqueous-alcoholic extract of the root culture of *Scutellaria baicalensis* were 30±0.2% ethanol at a raw material/extractant ratio of 1:86 at a temperature of 70±0.1°C for 6±0.1 hours. The scheme for the isolation and purification of trans-cinnamic acid obtained from the extract of the root culture of *Scutellaria baicalensis* is presented in Figure 1. The purity of the isolated trans-cinnamic acid was at least 95%.

**Figure 1.**
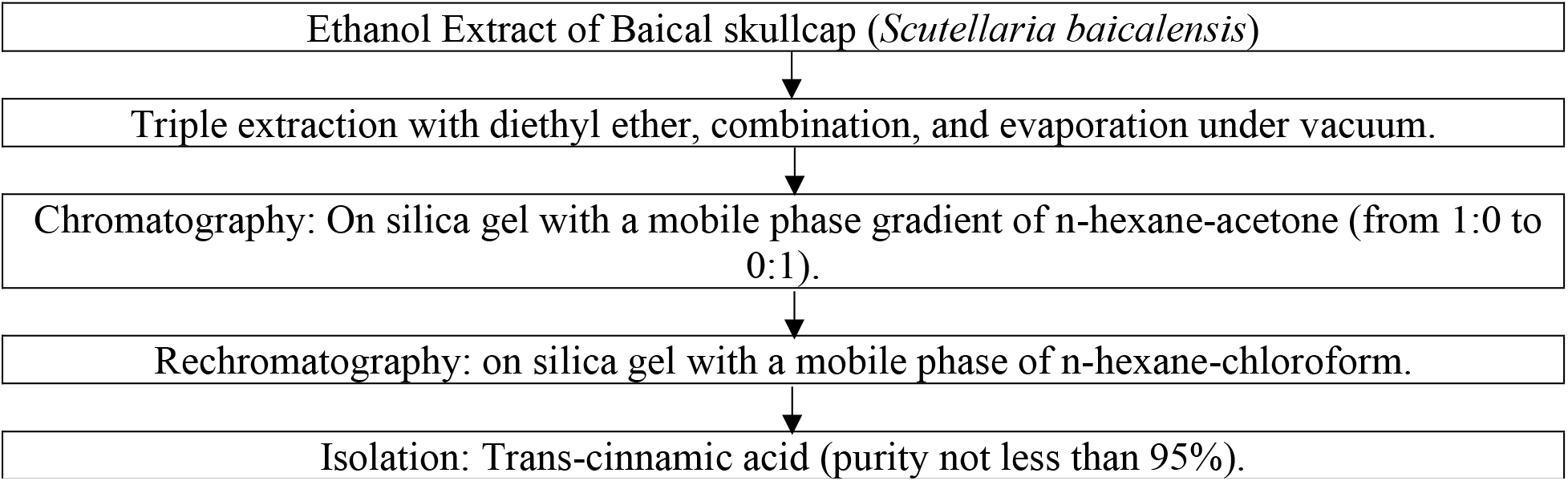
Scheme for the purification of trans-cinnamic acid obtained from the root culture extract of *Scutellaria baicalensis*.

The *in vivo* studies were conducted at LLC “Ifar.” The use of animals in this study and their housing conditions during the study were reviewed by the organization’s bioethics committee for compliance with the LLC “Ifar” Policy on Laboratory Animal Use and relevant regulatory documents governing work with laboratory animals [13–18].

To evaluate hypoglycemic activity, 50 male rats (Rattus sp.) with induced diabetes were used. Health status: pathogen-free, age at the start of the study – 12 weeks, body weight range at the start of the study – 219–272 g.

Males of this species were chosen because they do not have an estrous cycle, which affects susceptibility to etiological factors in females. This makes conducting the study on males preferable [13].

The study of the hypoglycemic activity of trans-cinnamic acid at doses of 50.0 mg/kg and mg/kg was carried out on model objects (male rats) with induced diabetes (by administering an alloxan solution). The design of the study on the hypoglycemic activity of trans-cinnamic acid is presented in Table 1.

**Table 1.** Design of the study on the hypoglycemic activity of trans-cinnamic acid.

| Group No. | Number of animals in the group | Modeling of diabetes | Substance administered, dose |
| --- | --- | --- | --- |
| 1 | 10 | - | - |
| 2 | 10 | + | Purified water |
| 3 | 10 | + | 50,0 mg/kg |
| 4 | 10 | + | 100,0 mg/kg |
| 5 | 10 | + | GLB, 5,0 mg/kg |

GLB – glyburide, antidiabetic substance

The study involved 50 rats, which were divided into 5 groups of 10 animals each [13]. Diabetes was induced in animals from groups 2–5, which had been previously deprived of food for 16 hours, by a single intraperitoneal injection of an alloxan solution at a dose of 150.0 mg/kg in a volume of 1.0 ml (conditions were experimentally pre-determined) [19]. The body weight of the animals and blood glucose concentration were measured before and 48 hours after the administration of alloxan to confirm the development of diabetes. Subsequently, the groups were formed only from animals with a blood glucose concentration of at least 11 mmol/L. To stabilize the condition of the animals, trans-cinnamic acid administration began 2 weeks later. Trans-cinnamic acid was given to the rats once daily for 7 days during the last week of the experiment. The body weight of fasting animals, blood glucose, and total cholesterol concentrations were assessed once a week for the 21 days of the experiment. Glibenclamide (GLB), a hypoglycemic agent, was used as a positive control at an effective dose of 5.0 mg/kg [20], administered intragastrically according to the same schedule as trans-cinnamic acid.

Comparison of feature values in several independent samples was performed according to SOP OFI-182. Outliers were evaluated according to GOST 16269-4-2017 [21] using Grubbs” test. Differences were considered statistically significant at p < 0.05.

## Results and Discussions

During the study, 17 rats died in the model of diabetes induced by a single intraperitoneal injection of alloxan solution at a dose of 150.0 mg/kg. As a result of the study, 1 mouse was removed from the hypercholesterolemia model due to signs of health impairment. The administration of trans-cinnamic acid at all doses did not cause severe health disturbances or mortality in the animals.

The average body weight data of rats after intraperitoneal administration of trans-cinnamic acid at doses of 50.0 and 100.0 mg/kg and GLB at a dose of 5.0 mg/kg are presented in Table 2.

**Table 2.** Average body weight data of rats after intraperitoneal administration of trans-cinnamic acid at doses of 50 and 100 mg/kg and GLB at a dose of 5 mg/kg.

| Time point | Body Weight, g |  |  |  |  |
| --- | --- | --- | --- | --- | --- |
|  | Intact animals | Purified water (control) | Trans-cinnamic acid |  | GLB |
|  |  |  | 50,0 mg/kg | 100,0 mg/kg | 5,0 mg/kg |
| Before injections | 242 ± 4 (n=10) | 243 ± 1 (n=80) |  |  |  |
| 48 hours after alloxan administration | 243 ± 4 (n=10) | 246 ± 4 (n=10) | 245 ± 3 (n=10) | 246 ± 5 (n=10) | 247 ± 3 (n=10) |
| 1 week after alloxan administration | 247 ± 4 (n=10) | 248 ± 2 (n=9) | 246 ± 2 (n=9) | 247 ± 4 (n=8) | 248 ± 5 (n=9) |
| 2 weeks after alloxan administration | 249 ± 3 (n=10) | 251 ± 4 (n=7) | 248 ± 3 (n=7) | 249 ± 4 (n=6) | 248 ± 4 (n=7) |
| 3 weeks after alloxan administration | 258 ± 3 (n=10) | 260 ± 4 (n=6) | 259 ± 2 (n=6) | 258 ± 3 (n=6) | 257 ± 4*<sup>#</sup> (n=6) |

«\*» denotes statistically significant differences from intact ( $p < 0.05$ ).
«#» denotes statistically significant differences from control ( $p < 0.05$ ).

The body weight dynamics of male rats in the diabetes model after administration of trans-cinnamic acid at doses of 50.0 and 100.0 mg/kg were comparable to those in the intact and control groups (p > 0.05). In rats with diabetes, receiving GLB at a dose of 5.0 mg/kg for 7 days, the body weight was comparable to the dynamics in the intact and control groups (p > 0.05). Thus, a single intraperitoneal injection of alloxan at a dose of 150.0 mg/kg and oral administration of trans-cinnamic acid at doses of 50.0 and 100.0 mg/kg did not affect the body weight dynamics of the experimental animals.

The average blood biochemistry data of rats after intraperitoneal administration of trans-cinnamic acid at doses of 50.0 and 100.0 mg/kg and GLB at a dose of 5.0 mg/kg are presented in Table 2.

**Table 2.** Average blood biochemistry parameters of rats after intraperitoneal administration of trans-cinnamic acid at doses of 50 and 100 mg/kg and GLB at a dose of 5 mg/kg.

| Time point | Intact animals | Purified water<br>(control) | Trans-coric acid |  | GLB |
| --- | --- | --- | --- | --- | --- |
|  |  |  | 50,0 mg/kg | 100,0 mg/kg | 5,0 mg/kg |
| Glucose, mmol/L |  |  |  |  |  |
| 48 hours after alloxan administration | 4,883 ± 0,301<br>(n=10) | 22,618 ± 0,792*<br>(n=10) | 22,872 ± 0,733*<br>(n=10) | 22,625 ± 1,280*<br>(n=10) | 21,475 ± 0,620*<br>(n=10) |
| 1 week after alloxan administration | 4,335 ± 0,227<br>(n=10) | 24,457 ± 0,700*<br>(n=9) | 24,565 ± 0,671*<br>(n=9) | 24,189 ± 1,106*<br>(n=8) | 25,134 ± 0,579*<br>(n=9) |
| 2 weeks after alloxan administration | 4,533 ± 0,285<br>(n=10) | 23,372 ± 0,721*<br>(n=7) | 24,433 ± 0,806*<br>(n=7) | 22,896 ± 0,436*<br>(n=6) | 22,861 ± 0,319*<br>(n=7) |
| 3 weeks after alloxan administration | 4,370 ± 0,378<br>(n=10) | 20,094 ± 0,398*<br>(n=6) | 20,372 ± 0,387*<br>(n=6) | 20,337 ± 0,350*<br>(n=6) | 13,022 ± 0,584*#<br>(n=6) |
| Cholesterol, mmol/L |  |  |  |  |  |
| 1 week after alloxan administration | 1,612 ± 0,063<br>(n=10) | 2,592 ± 0,090*<br>(n=9) | 2,514 ± 0,132*<br>(n=9) | 2,581 ± 0,099*<br>(n=8) | 2,583 ± 0,043*<br>(n=9) |
| 2 weeks after alloxan administration | 1,544 ± 0,088<br>(n=10) | 2,584 ± 0,074*<br>(n=7) | 2,411 ± 0,108*<br>(n=7) | 2,544 ± 0,089*<br>(n=6) | 2,554 ± 0,066*<br>(n=7) |
| 3 weeks after alloxan administration | 1,580 ± 0,074<br>(n=10) | 2,437 ± 0,073*<br>(n=6) | 2,499 ± 0,090*<br>(n=6) | 2,532 ± 0,063*<br>(n=6) | 1,927 ± 0,042*#<br>(n=6) |

«\*» – denotes statistically significant differences from intact ( $p < 0.05$ ).
«#» – denotes statistically significant differences from control ( $p < 0.05$ ).

Before the administration of alloxan, the glucose concentration in the serum of intact animals (5.264 ± 0.374 mmol/L, n=10) and the pool of animals for group formation (5.254 ± 0.163 mmol/L, n=80) corresponded to the norm. The values of these indicators did not differ from each other (p > 0.05).

Diabetes in rats developed 48 hours after a single intraperitoneal injection of alloxan at a dose of 150.0 mg/kg (p < 0.05), as evidenced by the increase in glucose and cholesterol concentrations in the serum of rats compared to the values in intact animals. The effect persisted in the experimental animals throughout the experiment (p < 0.05).

This allowed forming 10 individuals in each experimental group, however, by the end of the experiment, each group had 6 rats due to the toxicity of alloxan. The sample size is sufficient for statistical analysis and does not contradict ethical principles [22].

Oral administration of trans-cinnamic acid at doses of 50.0 and 100.0 mg/kg to experimental rats for 7 days did not lead to a decrease in glucose and cholesterol concentrations in the serum (p > 0.05). GLB at a dose of 5.0 mg/kg reduced the concentrations of glucose and cholesterol in the serum, but not to the level of intact animals (p < 0.05).

Thus, under these experimental conditions, trans-cinnamic acid at doses of 50.0 and 100.0 mg/kg did not exhibit hypoglycemic activity.

## Conclusion

The present study aimed to assess the hypoglycemic activity of trans-cinnamic acid extracted from the extract of *hairy rots Scutellaria baicalensis* in *in vivo* experiments. During the experiment, it was found that a single intraperitoneal injection of alloxan at a dose of 150.0 mg/kg and oral administration of trans-cinnamic acid at doses of 50.0 and 100.0 mg/kg did not affect the body weight dynamics of experimental animals with diabetes. Thus, under these experimental conditions, trans-cinnamic acid at doses of 50.0 and 100.0 mg/kg does not possess hypoglycemic activity. Therefore, dietary supplements containing these doses of trans-cinnamic acid are relatively safe, and their consumption is advisable.

## Funding

*The work was carried out within the framework of the state assignment on the topic “Development of biologically active additives consisting of metabolites of plant objects in vitro for protecting the population from premature aging” (project FZSR-2024-0008)*.

*The work was conducted using the equipment of the Center for Collective Use “Instrumental Methods of Analysis in the Field of Applied Biotechnology” at KemSU*.

## References

1. Polischuk, Yu. I. Analysis of the assortment of oral hypoglycemic medicinal products / Yu. I. Polischuk // Current Issues in Modern Medicine: Materials of the IV Far Eastern Medical Youth Forum, Khabarovsk, October 2–17, 2020. – Khabarovsk: Far Eastern State Medical University, 2020. – P. 287–288.

2. Severina, A.S. Place of dietary supplements in the treatment of diabetes mellitus / A.S. Severina, M.V. Shestakova // Diabetes mellitus. – 2007. – Vol. 10(2). – P. 76–79

3. Yakimova, T.V. Influence of extracts of medicinal plants on metabolic disorders in a model of diabetes mellitus and insulin resistance / T.V. Yakimova, O.N. Nasanova, A.I. Vengerovskiy // Bulletin of Siberian Medicine. – 2015. – No. 2. – P. 75–81

4. Dietary supplement use by US adults: data from the National Health and Nutrition Examination Survey, 1999-2000 / Radimer K., Bindewald B., Hughes J., Ervin B., et al. // Am J Epidemiol. – 2004. – V. 160(4). – Р. 339–49. doi: 10.1093/aje/kwh207

5. Jafarova, R.E. Study of preparations of some plant species to identify hypoglycemic action / R.E. Jafarova // Biomedicine. – 2007. – No. 4. – P. 27–32.

6. Yakimova, T.V. Influence of extracts of medicinal plants on metabolic disorders in a model of diabetes mellitus and insulin resistance / T.V. Yakimova, O.N. Nasanova, A.I. Vengerovskiy // Bulletin of Siberian Medicine, 2015, vol. 14, no. 2, pp. 75–81

7. Optimization of parameters for obtaining callus, suspension, and root cultures of meadowsweet (Filipendula ulmaria) to isolate the largest number of biologically active substances with geroprotective properties / S. Dyshlyuk, A. D. Vesnina, A. Y. Prosekov et al. // Brazilian journal of biology. – 2022. – V. 84. – P. 257074.

8. Antimicrobial and antioxidant activity of Panax ginseng and Hedysarum neglectum root crop extracts / L. S. Dyshlyuk, N. V. Fotina, I. S. Milentyeva [et al.] // Brazilian Journal of Biology. – 2024. – Vol. 84. – P. 256944

9. Study of the Effect of Baicalin from Scutellaria baicalensis on the Gastrointestinal Tract Normoflora and Helicobacter pylori. / A. Dmitrieva, I. Milentieva; A. Vesnina, A. Prosekov [et al.] // Int. J. Mol. Sci. – 2023. – Vol. 24. – P. 11906.

10. Biologically active compounds in Scutellaria baicalensis L. callus extract: Phytochemical analysis and isolation / I. S. Milentyeva, A. M. Fedorova, T. A. Larichev, O. G. Altshuler // Foods and Raw Materials. – 2023. – 11(1). – P.172–186

11. Research progress of active ingredients of Scutellaria baicalensis in the treatment of type 2 diabetes and its complications / W. Yingrui, L. Zheng, L. Guoyan, W. Hongjie // Biomed Pharmacother. – 2022. – Vol. 148. – P. 112690

12. Geroprotective activity of trans-cinnamic acid isolated from the Baikal skullcap (Scutellaria baicalensis) / A. M. Fedorova, L. S. Dyshlyuk, I. S. Milentyeva [et al.] // Food Processing: Techniques and Technology. – 2022. – Vol. 52, No. 3. – P. 582-591. – DOI 10.21603/2074-9414-2022-3-2388

13. Guide to Preclinical Studies of Medicinal Products. Part One / Ed. by A.N. Mironov. — Moscow: Grif and K, 2012. — 944 p.

14. GOST 33215—2014 Guide to the Maintenance and Care of Laboratory Animals. Rules for Equipment of Premises and Organization of Procedures — Moscow: Standartinform. 2019. — 12 p.

15. GOST 33216—2014 Guide to the Maintenance and Care of Laboratory Animals. Rules for the Maintenance and Care of Laboratory Rodents and Rabbits — Moscow: Standartinform. 2016. — 9 p.

16. GOST R ISO 10993-2—2009 Medical Devices. Biological Evaluation of Medical Devices. Part 2. Requirements for Animal Handling — Moscow: Standartinform. 2009. — 11 p.

17. SP 2.2.1.3218—14 Sanitary and Epidemiological Requirements for the Arrangement, Equipment, and Maintenance of Experimental-Biological Clinics (Vivariums) Approved by the Chief State Sanitary Doctor of the Russian Federation on August 29, 2014 No. 51.

18. Chlorogenic acid confers robust neuroprotection against arsenite toxicity in mice by reversing oxidative stress, inflammation, and apoptosis / D.M. Metwally, R.A. Alajmi, M.F. El-Khadragy [et al.] // Journal of functional foods. – 2020. – V. 75. – P. 104202. DOI:10.1016/j.jff.2020.104202

19. Ighodaro, O.M. Alloxan-induced diabetes, a common model for evaluating the glycemic-control potential of therapeutic compounds and plants extracts in experimental studies / O.M. Ighodaro, A.M. Adeosun, O.A. Akinloye // Medicina (Kaunas). — 2017. — Vol. 53, N 6. — p. 365–374

20. Rabbani, S.I. Protective role of glibenclamide against nicotinamide-streptozotocin induced nuclear damage in diabetic Wistar rats / S.I. Rabbani, K. Devi, S. Khanam // J. Pharmacol. Pharmacother. — 2010. — Vol. 1, N 1. — p. 18–23

21. GOST R ISO 16269-4-2017 Statistical Presentation of Data. Part 4: Detection and Treatment of Outliers. — Moscow: Standartinform. 2017. — 53 p

22. Russell W.M.N. The principles of humane experimental technique / W.M.N. Russell, R.L. Bunch — London: Methuen, 1959. — 258 p

